# Nano Colonies: Rearing honey bee queens and their offspring in small laboratory arenas

**DOI:** 10.1101/2024.01.20.576222

**Authors:** Zachary S. Lamas, Serhat Solmaz, Cory Stevens, Jason Bragg, Eugene V. Ryabov, Shayne Madella, Miguel Corona, Jay D. Evans

## Abstract

Honey bees create complex societies of self-organized individuals in intricate colonies. Studies of honey bees are carried out in both the field and the laboratory. However, field research is encumbered by the difficulties of making reliable observations and environmental confounders. Meanwhile, laboratory trials produce data that are not field realistic as they lack key characteristics of a natural colony. Additionally, advances in honey bee research have been hindered without reliable methodology to rear queens in the laboratory. Here we provide a new system to reliably produce queens and worker brood in the laboratory and describe how this system fits with artificial insemination of queens as a step towards a continuous self-contained source of bees. The process creates a bridge between field research and laboratory trials and provides a secure system for contagious or regulated elements while maintaining many of the intrinsic characteristics of a honey bee colony.

## Introduction

Honey bees are social organisms that are important to our economy by providing key pollination services for many food crops and producing honey and wax^1^. Ongoing research into honey bee health is integral for understanding factors that impact colony health. However, there is gap in research methodology that impedes advances; there is no reliable method to rear honey bee queens in the laboratory or to raise bee-reared progeny in the laboratory^2^.

Researchers study honey bees in both field and laboratory conditions. Traditional field trials provide real-world results and maintain all colony conditions, while laboratory trials lack key colony characteristics. However, field trials are costly, laborious, and vulnerable to environmental variation. Confounding variables are an inherent problem with field trials due to the difficulty in assessing and accounting for landscape effects including microclimate, pesticide exposure, and interactions with bees encountered during flights that span multiple kilometers^3^.

Laboratory trials offer a more uniform environment as the researcher can control all parameters in the experimental design. Arenas are typically made from clear plastic, allowing complete visibility to the researcher^4–6^. Due to the ease of setup and low cost, high numbers of replicates are more achievable and confounding variables are limited. Laboratory trials have created great advances in honey bee research, especially in regard to novel biossays^5,6^. However, laboratory trials exclude many components inherent to the colony environment. Specifically, laboratory arenas lack key colony characteristics such as brood, wax, a queen and associated pheromones, communication pathways and social behaviors that are inherent to field colonies. The absence of these key characteristics makes it difficult to extrapolate findings from the laboratory to the real world.

Current *in vitro* methodology allows researchers to rear brood artificially in the laboratory^7^. The methodology utilizes artificial feedings by a technician rather than brood care from an attending nurse bee. As a result, the hypopharyngeal glands, a gland unique to hymenopterans that generates a processed food source^8^, are completely bypassed. What is needed is a bridge between the real-world colony environment provided in field trials with the low cost and reliability of cage trials.

Here we describe one such bridge. We have developed a method to get a small number of worker bees to both rear queens in the confines of the laboratory and support a laying queen and raise her offspring. These laboratory colonies can be made with as few as 100 bees housed in a typical incubator. The colonies are referred to as queen-rearing or queen-right “nanos” or “nano-colonies”. They provide a next step in laboratory methodology to help bridge the complexities of a colony with the simplicity of the laboratory environment.

In this study we experimentally produced queens in laboratory settings. In several trials these queens were allowed to open mate, and we compared their mating success and morphology to traditionally produced sister queens. In another trial, we produced and instrumentally inseminated queens in the laboratory and then supported *in vivo* brood rearing without exposing these queens to the field. Finally, we experimentally produced brood through a reliable *in vivo* laboratory design.

## Methods

### Overview

#### Queen-Right Nanos

*Queen-right nanos consist of 100-240 worker bees and a queen. The hive is made from a modified plastic cup ventilated with #8 hardware cloth and noseum netting. A portion of empty, dark (seasoned in colonies after multiple rounds of worker bee production) brood comb serves as the brood area for the colony. Young worker bees are installed either soon after emergence or by being scooped from a brood frame*.

#### Queen-Rearing Nanos

*Queen-rearing nanos are queenless colonies of approximately 100 bees in modified plastic cages that will rear their own queen. The colonies are easy to establish and maintain in the laboratory. Establishment is completed over 2 steps. First, a strong queenless colony with ample provisions and young worker bees (a ‘cell builder’ colony) is made in the field. First instar larvae are grafted into plastic queen-rearing cages and placed into the cell builder for 12-24 hours. Next, nano colonies are made by securing an initiated queen cell into the nano, and then harvesting a small number of bees (approximately 100) from the cell builder and securing them inside the modified plastic cage. The nano colonies are then given feeders, sealed, and placed into an incubator*.

##### Making the Modified Cages

The cages are modified by removing the lower third of a 16 oz clear plastic cup, and then securing #8 hardware mesh to the bottom opening with hot glue. The lid of the cage is comprised of two materials: 4×4 inch piece of ‘noseeum’ mesh, and a lid to the plastic cup with the center cut out. A piece of paraffin wax honey comb (Betterbee, Inc.) is cut to fit into the center of the cup for the queen-rearing nanos. In contrast, empty, dark brood comb works best for the queen-right nanos because it allows for easy observation of eggs and young larva. The comb is placed in a −20 freezer for 10 minutes, and then cut into squares on a table saw to fit inside the modified plastic cups.

2 ml Eppendorf tubes are used as feeders for each colony type. For the sucrose and water feeders, a 2 mm hole is made in the bottom of the tube to allow bees to feed on the contents. Holes were made by perforating the tube with either a small brad nail or a 5/64 drill bit. A small cut in the noseeum lid is made with dissecting scissors and the feeder is then squeezed into this slit. The fabric will stretch as the tube is inserted. The lid of the tube is wider than the hole made by the bottom, and as such the feeder remains suspended in the lid. Pollen feeders were made by cutting 5 mm diameter opening in the bottom of an Eppendorf tube with a razor blade, and then packing the tube with either bee bread or pollen substitute (MegaBee) mixed according to the manufacturer’s directions. Pollen feeders were changed every other day. Each queen-rearing nano held 1 water feeder, 2 sucrose feeders (1:1 sucrose w/v) and 1 pollen feeder (Fig. 3). These feeders were changed daily. Seven ml of Supplementary diet (*see Supplementary Information* Table 1-2) is mixed with 43 ml of honey. The honey mixture is placed into feeders which are identical to the pollen feeders, and inserted into the cages from underneath so that the bees may access the honey similar to Shpigler et al^5^. A step by step guide to cage construction can be found in *Supplementary*.

##### Establishing queen rearing nano colonies

The queen-rearing nano colonies need two components to be established: a started queen cell and nestmate bees to complete queen rearing. First, queen cells are established by grafting 1^st^ instar worker larvae into plastic queen cups (JZ/BZ) and placing these into a traditional cell builder for 12-24 hours. During this time the nestmate bees of the cell builder will engorge the larva on a bed of royal jelly. We consider this a *started queen cell*. The queen cups are inspected 12-24 hours post insertion, gently rolled upwards and visually inspected for a small pool of jelly below a c-shaped larva. Empty cups or cups without a larva are not considered viable.

The second component is to populate the cages with worker bees who will attend and finish rearing queen cells. Workers can be acquired in one of two ways. A frame of emerging brood can be placed into an incubator and allowed to emerge, after which 100 newly emerged bees are placed inside a nano cup, provided a protein source, sucrose, and water, and then placed back into an incubator at 33°C for three days in order to mature. Conversely, workers can be obtained directly from the cell builder by shaking the nurse bees from a central frame of the cell builder into a plastic tub, and then scooping 1/3^rd^ cup of bees into a nano, and then quickly sealing it. For the first method, the queen cell is inserted into the cover of the nano for the bees to access it. For the second method, the started cell cup is directly pressed against the wax comb inside the nano, and then the nurse bees are added. We used a digital scale and measured 100-130 grams of bees to estimate the nurse bee population. We did this in sets of ten without any observable stress or loss to the queen cells.

##### Inspecting the nano colonies

Egg laying, brood development, queen cell development and nestmate mortality are all observable in queen-right or queen-rearing nano colonies. Although the plastic cup arena is inexpensive and easily procured, the concave nature of the cup does not allow for quantification of egg laying or brood development. Observers have some ability to view eggs and track larval development by tilting the cup while shining a light into the cage. We made reference cells prior to nano establishment by plunging a white-out applicator into a series of cells. During inspection these marked cells act as convenient reference points for egg laying and brood development in adjacent cells. The observer simply has to record the status of adjacent cells. An example of a typical queenright nano colony can be seen in Figure 1.

**Figure 1:**
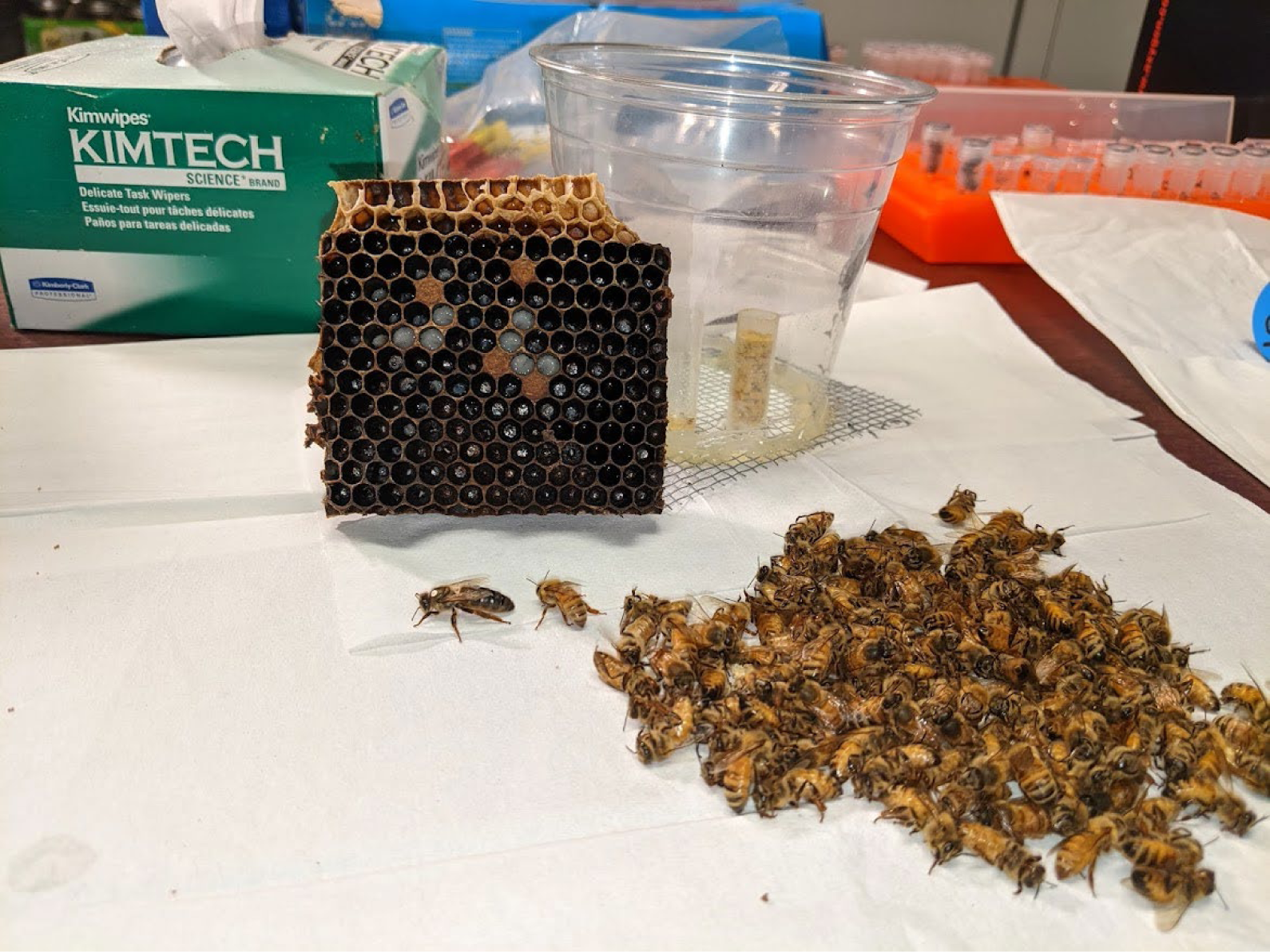
A typical queen-right nano 11 days after establishment. Center is one half of the brood area with capped and uncapped brood visible. Center and right, the adult bee population was anesthetized with carbon dioxide. The marked queen, and the entire worker population is presented.

**Figure 2:**
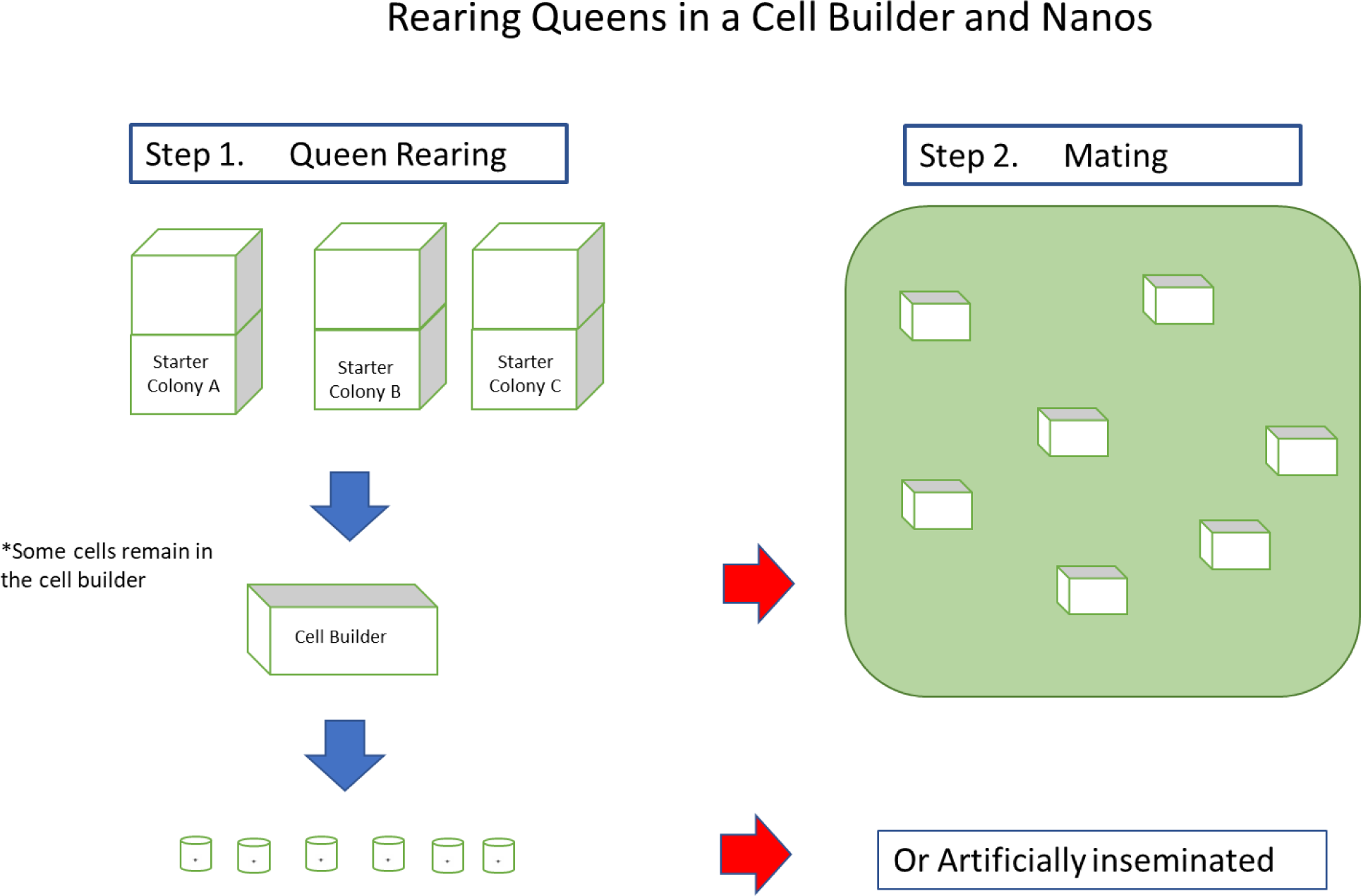
Step 1: Nurse bees are procured and placed into a cell builder. 1^st^ instar larvae are grafted into queen cups and placed inside the cell builder for 12-24 hours for initiation. Afterwards, queen cells are placed into nano colonies and given nest mate bees to continue queen rearing. Step 2: Mature queen cells (10 days after grafting) can be moved to traditional mating nucs for open mating or they can remain in the lab for instrumental insemination.

We used preliminary trials to test an arbitrary range of how many adult worker bees were needed to establish an artificial colony. Colonies with both low and high ranges of adult bees (100 – 240) successfully reared brood.

Bees to establish nanos can be collected either from emerging brood frames or directly from the brood frames in a field colony. Frames of emerging brood from healthy colonies exhibiting no covert signs of disease are placed into an incubator at 33°C and 60% humidity in a frame holder. Bees are collected within 24 hours of emergence, and placed into a nano setup. For the latter, when collecting bees directly form a colony, bees collected directly from colony brood frames are placed into the nano colony. Queens can be installed prior to or soon after worker addition. Colonies are then maintained in an incubator at 33°C.

**Figure 3.**
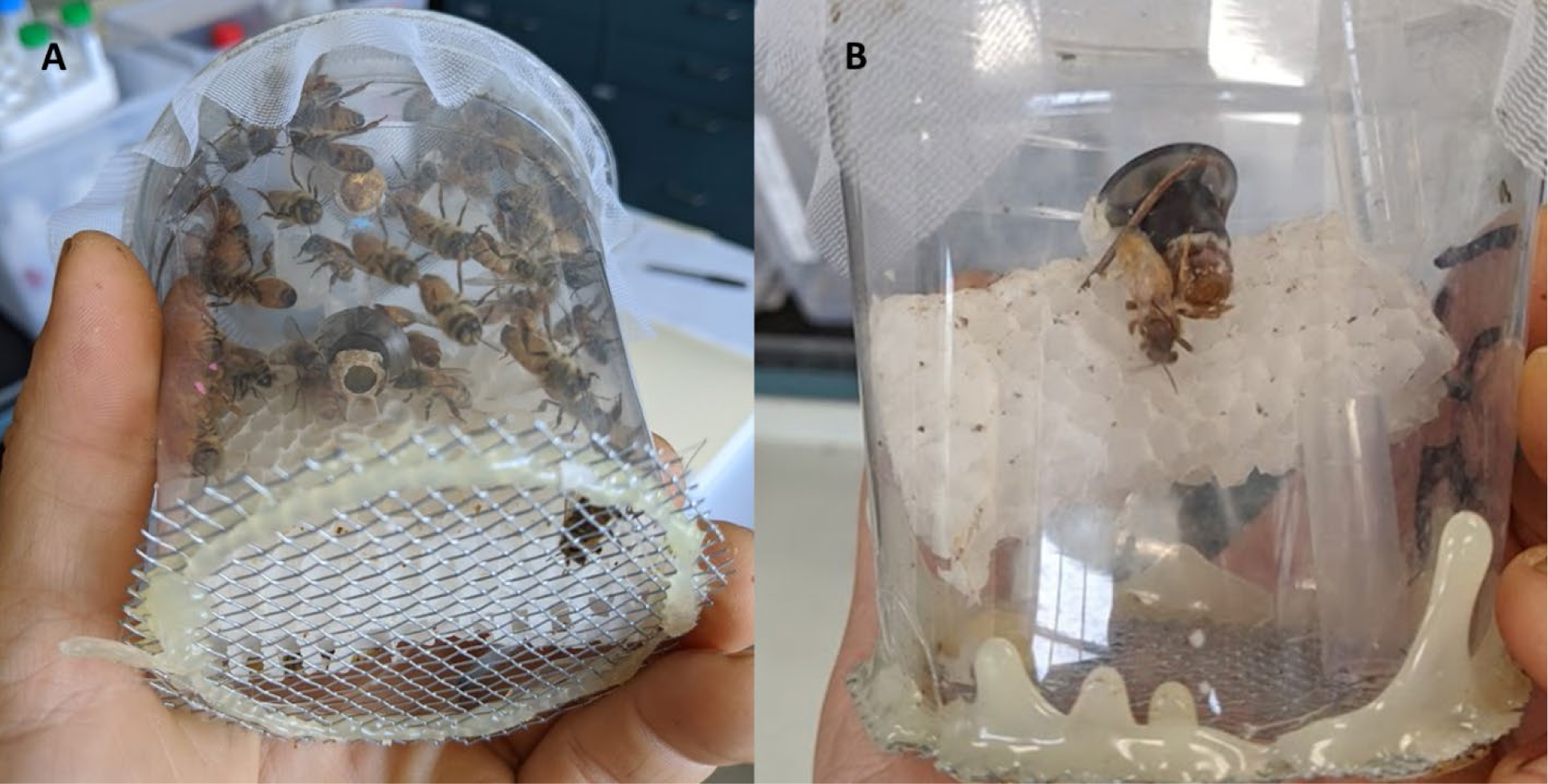
The arena depicted above produced virgin queens using pollen substitute, sucrose and parafilm comb, and approximately 130 worker bees in the arena. **(A)** A queen-rearing nano colony with open queen cell and larva viewable from the underside of the arena. The queen cell depicted is 24 hours after insertion into the nano colony. **(B)** A virgin queen newly emerged from her pupal cell. Not seen are the worker bees which were anesthetized with carbon dioxide and removed prior to queen emergence.

##### Comparing mating success of queens reared in traditional cell builders to laboratory reared queens

In two trials, queens were reared in the laboratory in parallel with sister reared queens in a traditional cell builder. The two groups were then compared for mating success. During the second trial, additional treatments within the nano group were added. We tested if queens could be reared not just on bee bread, but also artificial pollen substitute. We created a third group testing if the amount of time a queen cell was started in a traditional cell builder could be reduced to as little as 12 hours before being transferred into a nano colony for completion. These additional groups had few replicates in each group (N = 6).

##### Instrumental insemination of lab reared queens

We produced queens in the laboratory in two different trials, and then instrumentally inseminated the queens. The queens were re-introduced into nano colonies to lay.

## Results

### Queen-right nanos

In two different trials, queen-right nanos were established (N = 24), and maintained for 18 days. All colonies supported a laying queen and produced brood during this time period. Brood progression was visually analyzed on day 18 when the nanos were deconstructed. The majority of nanos had developed brood, 23/24 (95.8%). Of these, 13 had capped brood, with an additional 3 having late stage, L5, larvae, but no capped brood. Manual inspection of the worker brood revealed typical worker pupae, which can be seen in figure 1. No experimental variables were tested in these trials, but rather the experiments were carried out as a proof of concept if a small number of bees could rear brood in laboratory settings. Queens were overnight mailed to the Queen and Disease Clinic at North Carolina State University Extension for morphometric, sperm viability and sperm quantity analysis.

### Queen-rearing nanos

#### Comparing mating success of queens reared in traditional cell builders to laboratory reared queens

There was no difference in mating success between the two groups in the first trial: 6/14 (42.9%) of the queens in each group successfully mated. More queens successfully mated in the cell builder group (7/10, 70%) than any of the nano groups in the second trial (37.5% across all nano groups, figure 4).

**Figure 4:**
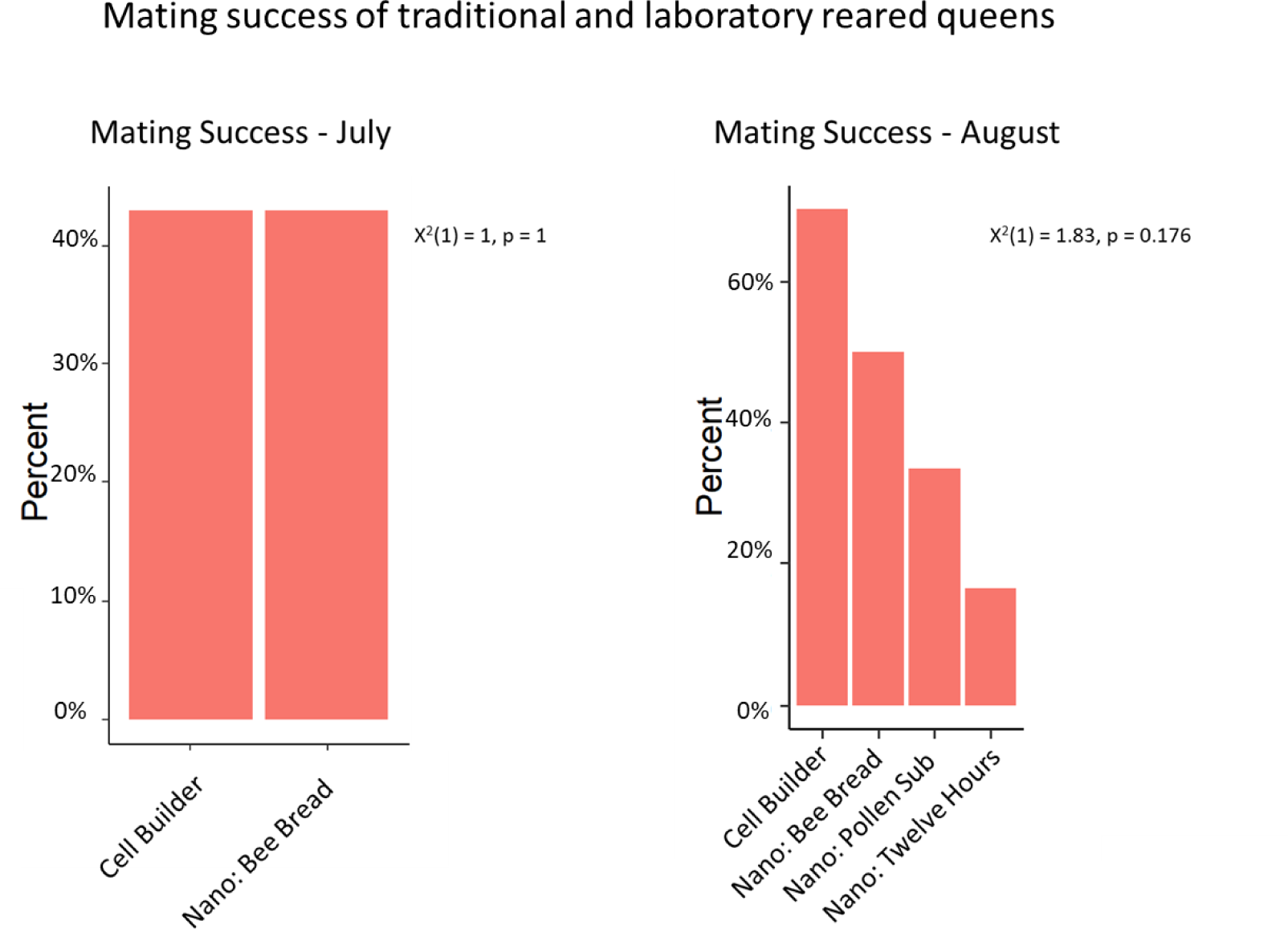
Mating success was measured between virgin queens produced using a traditional cell builder and virgin queens reared in the laboratory. There was no difference in mating success during the July trial. More queens by a proportion successfully mated in the traditional cell builder trial (70%) than any other laboratory group (17% - 50%) during the August trial. The differences were not statistically significant, X^2^(1) = 1.83, p = 0.176.

#### Morphometrics of queens reared in a traditional cell builder to laboratory reared queens

There was no difference between the body weights of queens between the cell builder (M = 192.77, SD = 9.73) or nano groups (M = 184.08, SD = 11.2), t(14.23)=1.78, p = 0.097. Nor was there a difference in sperm viability between the two groups: cell builder (M = 0.7957, SD = 0.0709) and nano (M = 0.8408, SD = 0.0464), t(9.064)=-1.51, p = 0.166. There was also no difference in total sperm stored in the spermathecae of mated queens regardless of whether they were reared in a traditional cell builder (M = 4.24, SD = 2.29) or a nano colony (M = 5.5, SD = 1.42), t(8.76)-1.32, p =0.22. (figure 5)

**Figure 5.**
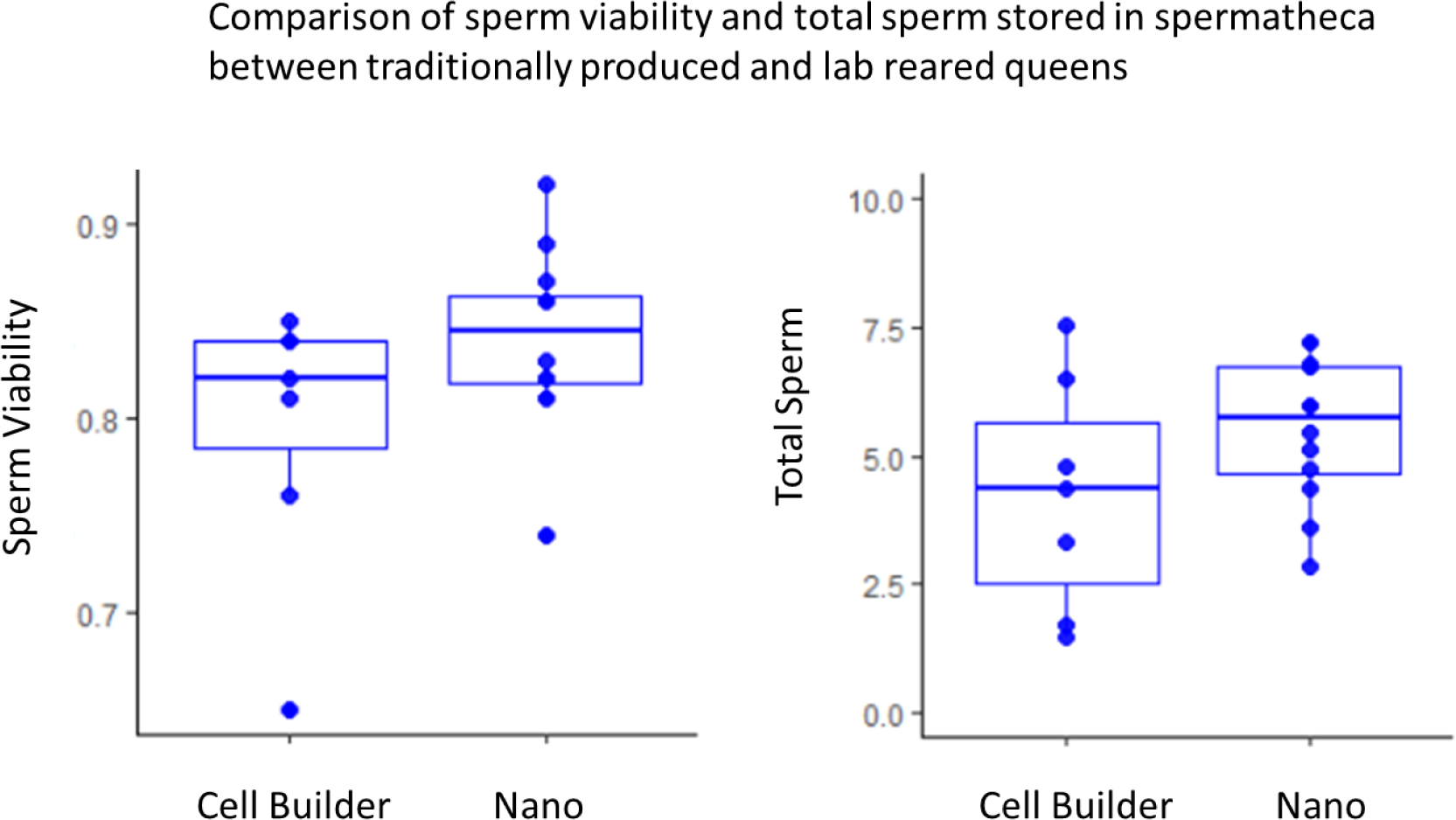
There was no significant difference between groups when analyzed for sperm viability, t(9.064)=-1.51, p = 0.166, and total sperm stored in queen spermatheca’s, t(14.23)=1.78, p = 0.097.

#### Instrumental insemination of laboratory-reared queens

Few queens successfully laid eggs, and fewer reared worker brood. In the first trial, queens did not immediately begin egg laying after initial carbon dioxide (CO_2_) exposure. Queens were anesthetized with CO_2_ a second time after insemination. Egg laying began 2 weeks after insemination. Nine queens began laying (9/23, 39.1%), but none of the colonies successfully reared late-stage larva. We did not confirm if the queens were laying fertilized eggs. In trial 2, queens were also anesthetized for a second time after insemination with CO_2_. Eight queens successfully laid eggs (8/12, 75%); Three colonies reared late-stage worker larva, and one nano colony, headed by a lab produced, instrumentally inseminated queen, successfully reared emerging adult worker bees.

## Discussion

Social insects create complex societies which can be difficult to study in field environments. Current laboratory systems simplify these societies into small groups while losing key characteristics. Here we provide a new laboratory method that utilizes a small number of bees to rear queens or worker brood while maintaining many colony characteristics. This method advances honey bee research by building a bridge between the complexities of field trials and the ease of use of laboratory research.

Our nano colonies bring more features inherent in honey bee colonies directly into the laboratory. Our colonies have laying queens, brood and honey comb. We hypothesize our setup could be useful studying exposure to disease or chemical stress on worker physiology. The absence of a queen or brood can affect worker bee physiology, limiting the relevance of current arenas. Our methodology provides a bridge between traditional field trials and the laboratory, and could be employed to advance other areas in honey bee research such as studying queen development, modified viruses, transgenic bees, or studying principle drivers of honey bee health.

Using a nano colony, researchers can study queen development and colony establishment in a closed environment and under controlled conditions. These queens can then either be returned to the field for natural mating or instrumentally inseminated, and then returned to a new nano colony setup to begin egg laying. We were able to produce queens after nominal larval development in the field, with a small number of nurse bees. Queens were successfully produced with either bee bread harvested from colonies or pollen substitute. We successfully reared and instrumentally inseminated queens in the laboratory, representing a closed loop methodology of queen to brood production, and brings a valuable toolset to the researcher. Queen right nano colonies were also used to develop worker brood within the confines of a laboratory incubator. A relatively small number of approximately 100 (100 – 150) bees supported a laying queen and reared her hatched eggs to the final stage of development as healthy pupae

Rearing queens and brood in the laboratory advances honey bee research by bridging the gap between field and laboratory trials. The method can be developed so that all components of each nano colony can be sampled and then examined against each other in a simple, closed system. In this way we may be able to parse out how individual drivers of honey bee health are impacting each component of a honey bee colony^9^. Variation can be further reduced by using pools of homogeneous bees. Researchers could potentially use this methodology to study the impact of one variable at a time on queen development or in a factorial design studying multiple variables at a time.

As is, this system can provide colony-like conditions into the laboratory, offering a closed system in which queens, workers, and developing bees can be observed and readily harvested for study. Nevertheless, this method can be further optimized. We did not test the resiliency of these laboratory colonies, and more work can determine the long term fitness of queens and worker bees reared in this way with different microbiome and nutritional provisions. We utilized inexpensive and widely available 16-ounce clear plastic cups to construct our colonies. This convex cup limits clear observation of developing brood but a flat design would allow for observation and quantification of brood development. Finally, it would be ideal to initiate queen rearing directly from eggs, so that their entire development could be controlled. We anticipate these are all solvable issues.

## Acknowledgements

Cory Stevens is an independent researcher at Stevens Bee Co., Bloomfield, MO, USA

Jason Bragg is an independent researcher at New River Honey Bees in Calvin, WV, USA. Queen morphometrics and sperm viability were provided by the Queen and Disease Clinic through North Carolina State Extension

## Funding

Funding was provided by the New Hampshire Beekeepers Association, the Cosmos Club Foundation, the Nansemond Beekeepers, and the Montgomery County Beekeepers.

## Step by step guide to constructing nano cages (1/5)

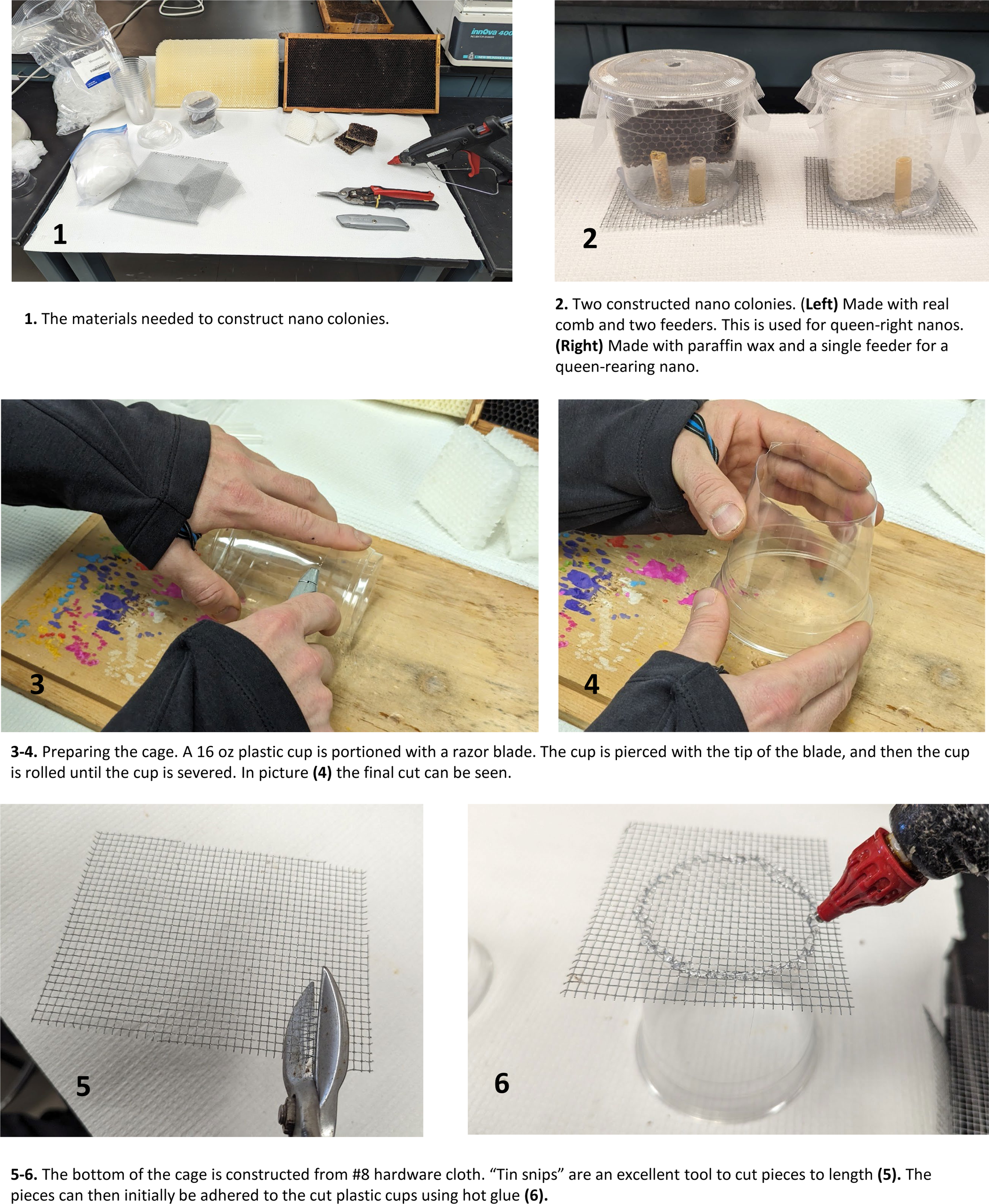

## Step by step guide to constructing nano cages (2/5)

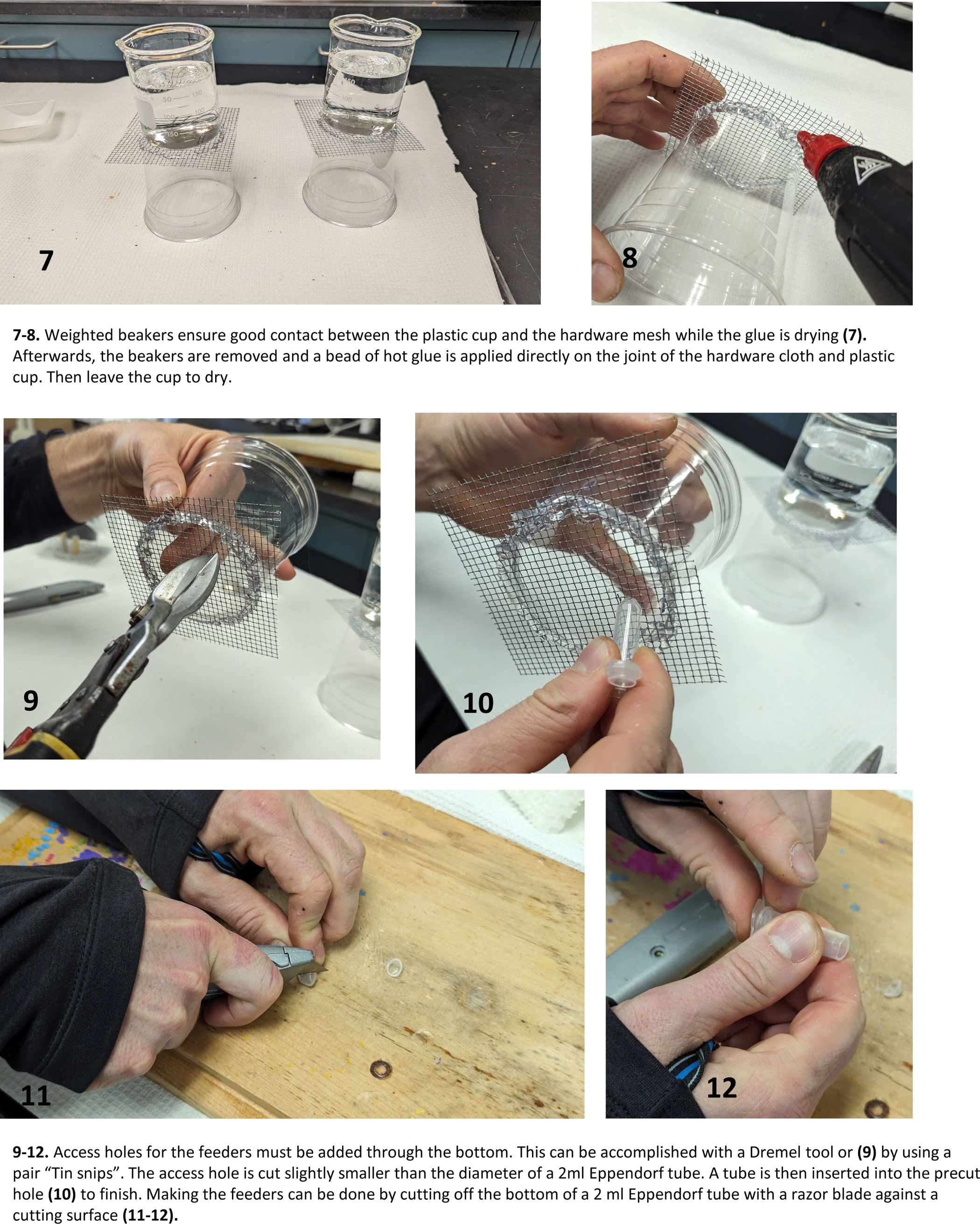

## Step by step guide to constructing nano cages (3/5)

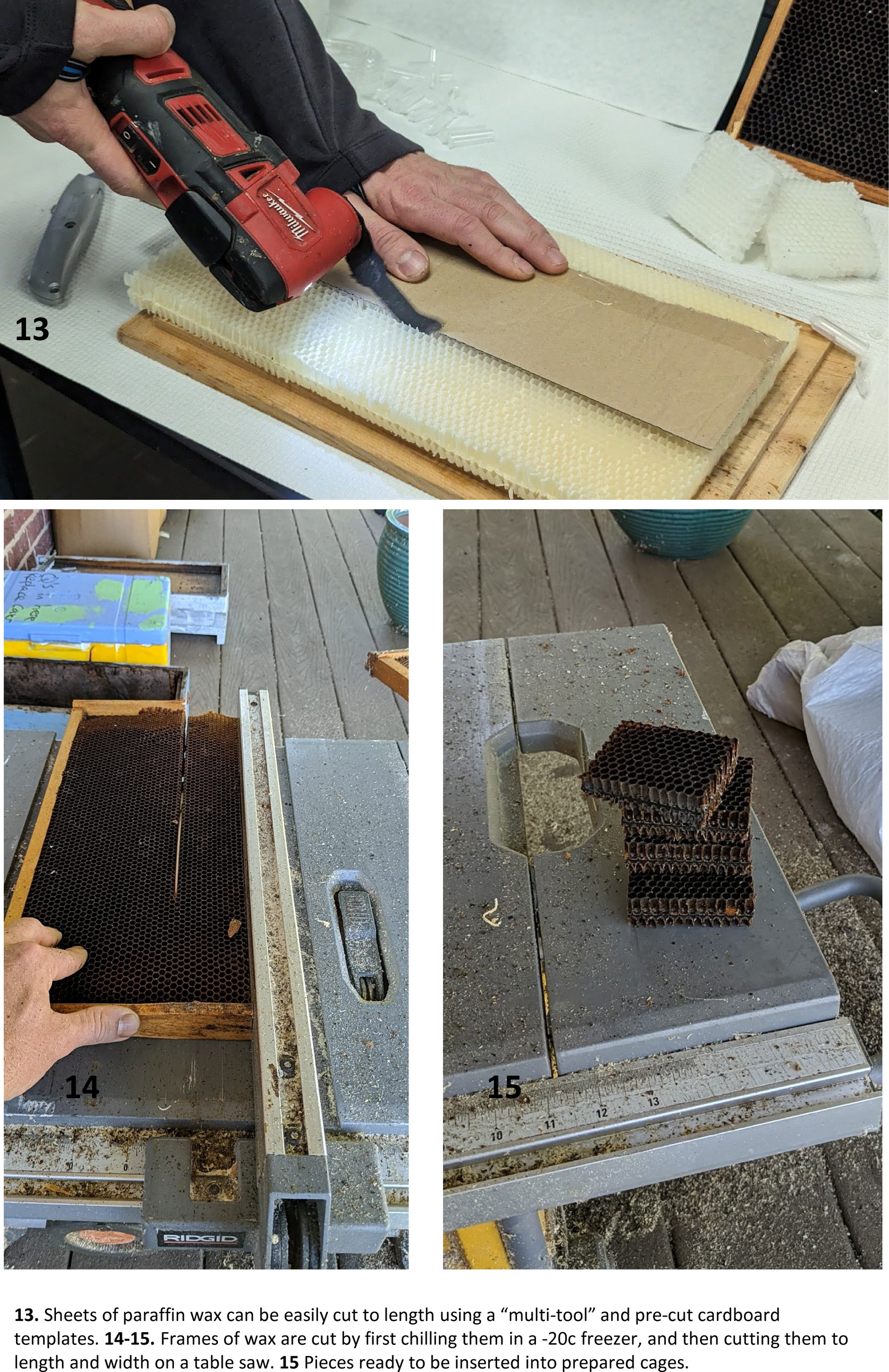

## Step by step guide to constructing nano cages (4/5)

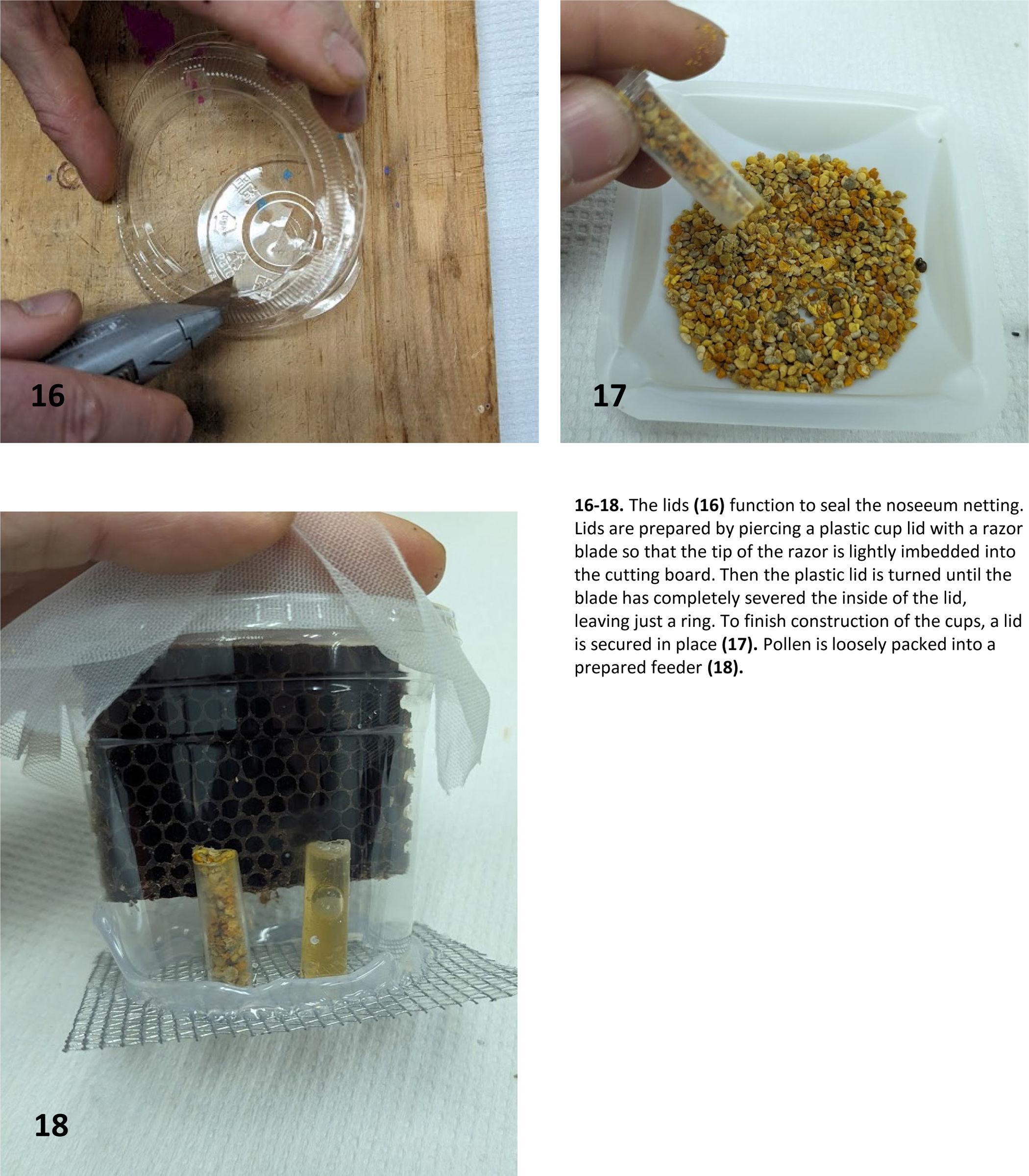

## Step by step guide to constructing nano cages (5/5)

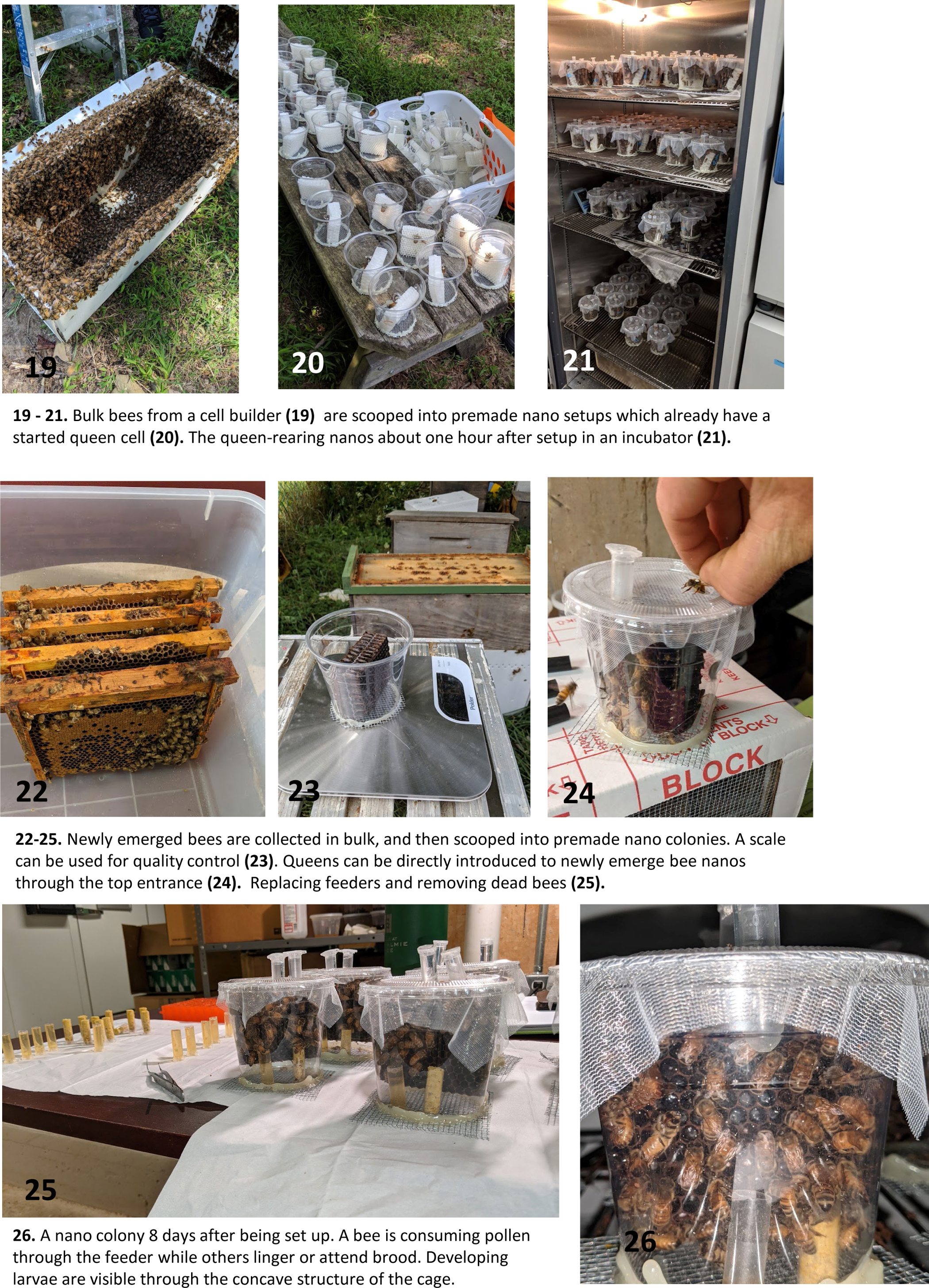

## Notes

### Competing Interest Statement

The authors have declared no competing interest.

## References

1 Khalifa, S. A. et al. Overview of bee pollination and its economic value for crop production. Insects 12, 688 (2021).

2 McAfee, A., Pettis, J. S., Tarpy, D. R. & Foster, L. J. Feminizer and doublesex knock-outs cause honey bees to switch sexes. PLOS Biology 17, e3000256, doi:10.1371/journal.pbio.3000256 (2019).

3 Corona, M., Branchiccela, B., Madella, S., Chen, Y. & Evans, J. Decoupling the effects of nutrition, age and behavioral caste on honey bee physiology and immunity. BioRxiv, 667931 (2019).

4 Evans, J. D., Chen, Y. P., Prisco, G. d., Pettis, J. & Williams, V. Bee cups: single-use cages for honey bee experiments. Journal of Apicultural Research 48, 300–302, doi:10.1080/00218839.2009.11101548 (2009).

5 Shpigler, H. Y. & Robinson, G. E. Laboratory Assay of Brood Care for Quantitative Analyses of Individual Differences in Honey Bee (Apis mellifera) Affiliative Behavior. PLoS One 10, e0143183, doi:10.1371/journal.pone.0143183 (2015).

6 Fine, J. D. et al. Quantifying the effects of pollen nutrition on honey bee queen egg laying with a new laboratory system. PloS one 13, e0203444–e0203444, doi:10.1371/journal.pone.0203444 (2018).

7 Schmehl, D. R., Tomé, H. V. V., Mortensen, A. N., Martins, G. F. & Ellis, J. D. Protocol for the in vitro rearing of honey bee (Apis mellifera L.) workers. Journal of Apicultural Research 55, 113–129, doi:10.1080/00218839.2016.1203530 (2016).

8 Snodgrass, R. E. Anatomy of the honey bee. (Cornell University Press, 1956).

9 Steinhauer, N. et al. Drivers of colony losses. Current opinion in insect science 26, 142–148 (2018).

